# Testing a Method for Quantifying Microplastic Content in *Mytilus californianus*

**DOI:** 10.1101/2024.03.04.583441

**Authors:** Ajeetesh S. Sidhu, William Sun, Mariam B. Helal, Aryaman Gupta, Aarsh Mittal, Serena Ramanathan, Manu Thakur, Bhoomi Jain, Dylan Wang, Andrew Benson, Connor T. Adams

## Abstract

With the proliferation of microplastics within the environment, there has been an increased need to quantify them within organic tissues of aquatic species. Several studies have examined the potential role of mussels in filtering these contaminants, prompting interest in characterizing uptake and retention of microplastics in these species. There are a variety of methods to isolate and quantify microplastics, including density separation, enzymatic digestion, and filtration. The applicability of a combined digestion and ultrasonic bath methodology in isolating microplastics from the California mussel (*Mytilus californianus Conrad, 1837*) was investigated. Microplastics were added to 20-gallon tanks at 0 g (control) 8 mg. Mussel tissues from each tank were digested using potassium hydroxide solutions. These digestions were subjected to vacuum filtration and a sonication bath, then treated with a mixture of sodium hypochlorite and nitric acid. It was found that this method successfully removed organic tissue from the mussels while leaving dyed microplastics intact. The latter were quantified using fluorescent microscopy. This proof-of-principle experiment indicates that this technique can be applied to further studies on environmental impact and retention of microplastics in mussels and may also have utility for other shellfish.

## Introduction

Microplastics (MPs) are fragments of plastics ranging in size from 10 μm to 5 mm that have become some of the world’s most prevalent pollutants due to their mobile and microscopic qualities. They have even been detected in human blood and placentas (Ragusa et al. 2021) (Wright, Kelly 2017) (Leslie et al. 2022) (Vandenberg et al. 2017). In addition, MPs are ubiquitous throughout aquatic environments (Wright et al. 2013). Alfaro-Núñez et al. (2021) found that 100% of organisms collected in the Tropical Eastern Pacific and the Galápagos Archipelago contained some level of MPs. These pollutants have spread across marine environments as ocean currents, zooplankton, and input from runoff drive the proliferation of MPs globally (Kvale et al. 2020). There are an estimated 24.4 trillion particles in the upper oceans (Isobe et al. 2021). According to one projection, plastics are expected to enter oceans at a rate of 9.6 to 48.8 particles per cubic meter by 2100, reflecting a 50-fold increase from today’s levels (Everaert 2018). Even more remote and fragile marine ecosystems, such as those found in the Antarctic and coral reefs, have begun to encounter MPs as well (Horton and Barnes 2020). Some of the most common types of plastics found in the ocean are polyethylene (PE), polypropylene (PP), and polystyrene (PS) (Shwarz et al. 2019); however, it is important to consider the prevalence and biological effects of other types of plastics as well.

Microplastics have been categorized as ecotoxicological pollutants with numerous known or possible negative effects on marine organisms, such as: tissue damage through oxidative stress, disruption of reproductive function, and serving as carriers of other toxic pollutants (Amobonye et al. 2021) (Bakir et al. 2016). Although research is still in its early stages, studies have indicated that MPs may also have an impact on human health. This includes possible adverse effects upon the endocrine, respiratory, reproductive, and immune systems (Bouwmeester et al. 2015) (Wright, Kelly 2017) (Vandenberg et al. 2017) (Leslie et al.2022) (Lee 2023). Due to the potential negative effects on marine organisms and ecological systems, and in light of the large amounts of MP present globally, it is important to validate MP quantitation techniques and tissue uptake measurements in shellfish and other species.

Mussels have emerged as a common organism to quantify MP content during ecological studies, since they are filter feeders and likely susceptible to bioaccumulation of MPs (Cauwenberghe and Janssen 2014). The filter-feeding capabilities of mussels allows them to extract MPs from the aquatic environment and store them within their own digestive tissues or feces (Fernández and Albentosa 2019) (Gonçalves 2019). Moreover, mussels are found globally (Böhm et al., 2021) and thus might be an interesting model to examine MP dynamics in multiple geographical sites and environments worldwide. Furthermore, mussels are a potential MP carrier into the human food supply (Grienke 2014). An emerging concept is use of mussels for ocean MP “clean up” (Al-Thawadi 2020) to complement other strategies such as large-scale floating barriers, use of robotics (Shmaltz 2020), and bacteria that capture and break down MPs (Sheth et al. 2019).

Despite multiple studies exploring the relationship between mussels and MPs, the degree to which tissue MPs are retained with various exposure levels remains to be fully elaborated. To support such studies, methods for MP measurements are required. In this study, a method for MP quantitation in mussel tissues was established by exposing California mussels (*Mytilus californianus*) to varying levels of MPs. The California mussel is an extremely prolific species that is commonly found along the Eastern Pacific Coast from Southern California to Alaska and considered to be a keystone species (Beauchamp and Gowing 1982). These mussels are bivalve feeders capable of trapping MPs of any size within their gills. Filtered particles are then sorted and eventually passed into the mouth, where they are digested and excreted (Gonçalves 2019) (Whedon 1938). California Mussels can filter 1.8-2.8 liters of water an hour (Bayne 1976) allowing mussel beds to filter large volumes of water within any given region. Past studies have found blue mussels, which are related to the California mussel, to have some adverse effects from exposure to MPs, such as a decreased clearance rate (volume of water filtered by an individual mussel over time) and perturbations in energy metabolism (Hamm and Lenz 2021). The dynamics of MP in *Mytilus californianus* and application of MP quantitation methods are lacking in this species. To this end, a proof-of-principle study was conducted in which tissue MP levels in mussels were determined through approximately weekly tissue digestion and filtrations over several months. The MPs were then analyzed quantitatively using Fourier transform infrared (FTIR) spectroscopy. These studies established methods to quantify the MP net retention of *Mytilus californianus*.

## Methods

### Mussel Collection and Care

Samples of *Mytilus californianus* were collected between 17 July 2022 to 31 July 2022 from Maverick’s beach (37.4996968909796 N, 122.49682473372347 W), an unprotected natural habitat within Half Moon Bay, California. These mussels were then organized into two 20-gallon (75.7 L) tanks with approximately 15 mussels per tank. The water was kept at 35 ppt salinity and at room temperature (20°C). Biweekly, 4 gallons (15.1 liters) of water were changed, with the amount of aquarium salt and Seachem Prime Aquarium Water Conditioner (Seachem Laboratories, Madison, GA) (0.1 mL conditioner/1 L water) kept constant to avoid shocking the mussels. In addition, on a regular basis the sides of the tanks were scraped to remove algae, tanks were regularly checked for dead mussels, and cleaned of large debris. Mussels were fed weekly with Seachem Laboratories Reef ZooPlankton store-bought feed and tank filters were cleaned on a biweekly basis.

### Addition and Formation of Microplastics

Microplastics were taken from a mixture made from equal masses of 6 types of common plastics: Plastic Number 1-Polyethylene Terephthalate (PETE), Plastic Number 2-High Density Polyethylene (HDPE), Plastic Number 3-Polyvinyl Chloride (PVC), Plastic Number 4-Low Density Polyethylene (LDPE), Plastic Number 5-Polypropylene (PP), and Plastic Number 6-Polystyrene (PS). Microplastics were created using an orbital sander to grind down typical consumer plastic, including plastic utensils (PP & PS), chip bags (PP & LDPE), food wrappers (LDPE), milk jugs (HDPE), Falcon tubes (PP), and water bottles (PETE & HDPE). Using an analytical balance, equal amounts of each type of microplastic were measured to create the final mix. Once weighed, MPs were placed into centrifuge tubes and treated with Nile Red Dye (1 μg/mL) in methanol (“Nile Red for Microplastics” solution, Crime Scene, Phoenix, AZ, USA). Enough was used to completely cover the sample (minimum 4 drops). Each sample was vortexed at moderate speed for 30 seconds. The sample was then allowed to sit for 30 minutes before the MP suspension was added to its respective tank. To treat the mussels, 0 mg of MP mix was added to the control tank and 8 mg to the MP tank.. Microplastics in the MP tank were maintained when water was changed, with 8 mg of MP mix added to the 4 gallons (15.1 L) of replacement water.

### Collection and Digestion of Tissues

A mussel was collected from each tank approximately week 1 (day 7), week 3 (day 19), week 5 (days 37 and 38), week 6 (day 44), and week 8 (day 58). Using a caliper and electronic balance, the wet weight of the mussel with the shell, its length and width were measured. Using a putty knife and disposable scalpel, each mussel was cracked open and the tissue and liquids inside were scraped and poured into a 50 mL conical tube (Falcon, Corning); wet tissue masses were calculated from tube weight measurements taken empty and then following addition of tissue. To digest the tissue, a 10% potassium hydroxide (KOH) solution in water was utilized, which breaks down organic tissue without dissolving polymers. KOH was chosen as the primary digestion chemical due to past studies finding it to cause no loss of MPs in biological matrices, with a 100% recovery rate (Dehaut et al. 2016). The KOH solution was poured into each tube so that the solution covered up to approximately twice the amount of tissue present within the tube. The tube was then stored in an incubator at 37°C for at least 72 hr without mixing and then transferred to a 4°C freezer. If the sample showed incomplete digestion after 72 hr it would be incubated further until all organic material was completely dissolved.

### Filtration/Sonication Process

To isolate and quantify the MPs, several steps were taken to remove tissues and organic materials. First, the digestion was combined with deionized water in a 3:1 ratio by adding water directly to the sample tube, then mixed. In a chemical fume hood, the digestate was then filtered through a plain white cellulose acetate membrane filter (0.45 μm, 47 mm; Advantech Manufacturing, Mentor, OH, USA) using a Buchner funnel. Deionized water was used to rinse any remaining digestate from the sample tube onto the Buchner funnel. A secondary digestion solution was prepared by combining 3 mL of nitric acid (HNO_3_), 9.25 mL of 1.24 M sodium hypochlorite (NaClO), and 20.75 mL of deionized water to obtain a final concentration of 0.3185 M for the NaClO. These chemicals were chosen for the wash due to their proven effectiveness at removing biological matter, as well as previous studies finding that short-term exposure to NaClO and HNO_3_ will not degrade microplastics (Collard et al. 2015). With the vacuum turned off, the secondary digestion solution (33 mL) was slowly added to the Buchner funnel and allowed to sit for 5 minutes before being vacuum filtered. The filter membrane was then carefully removed with forceps and placed into a 50 mL Falcon tube with enough methanol to fully cover the filter. The tube was sonicated in an ultrasonic bath at 50 hertz for 5 minutes. To store and evaporate the membrane and methanol, the methanol and filter were transferred into a plastic 60 mm petri dish, MP side up on the filter. Extra methanol was used to rinse the tube, ensuring the remaining MPs were rinsed onto the filter paper. The petri dishes were then placed into an incubator at 37°C to allow them to evaporate. After evaporation, each dish was rinsed with enough hydrogen peroxide to barely cover the filter to dissolve any leftover organic material, followed by another round of evaporation.

### Microscopy

The petri dish with filter paper (MP side facing up) was placed on the sample platform of a Zeiss AxioVert 200 inverted fluorescence microscope. To quantify MPs, first a picture was taken of the filter in the middle of the petri dish at 150X magnification, and the number of microplastics in the 0.3 mm^2^ area was manually counted through image analysis. This step was repeated for the left, right, top, and bottom of the petri dish. After all MPs in these zones were counted, an average was found to estimate the mean of the microplastics particles per 0.3 mm^2^. This was used to estimate the total MP within the whole dish which had a total area of 1450 mm^2^.

## Results

The mean (± SD) wet weight per mussel analyzed for MP content was 29.1459 ± 14.2088 g. Fluorescence microscopy indicated successful isolation of MPs following the digestion and filtration method approaches used herein (**Figures 1A-1C**). As illustrated in the figures, control samples without the addition of dyed MPs did not reveal false positives, while dyed sample fluorescence was great enough to clearly count individual MPs in both low and high tanks. All images have been made available in Appendix A.

**Figure 1A:**
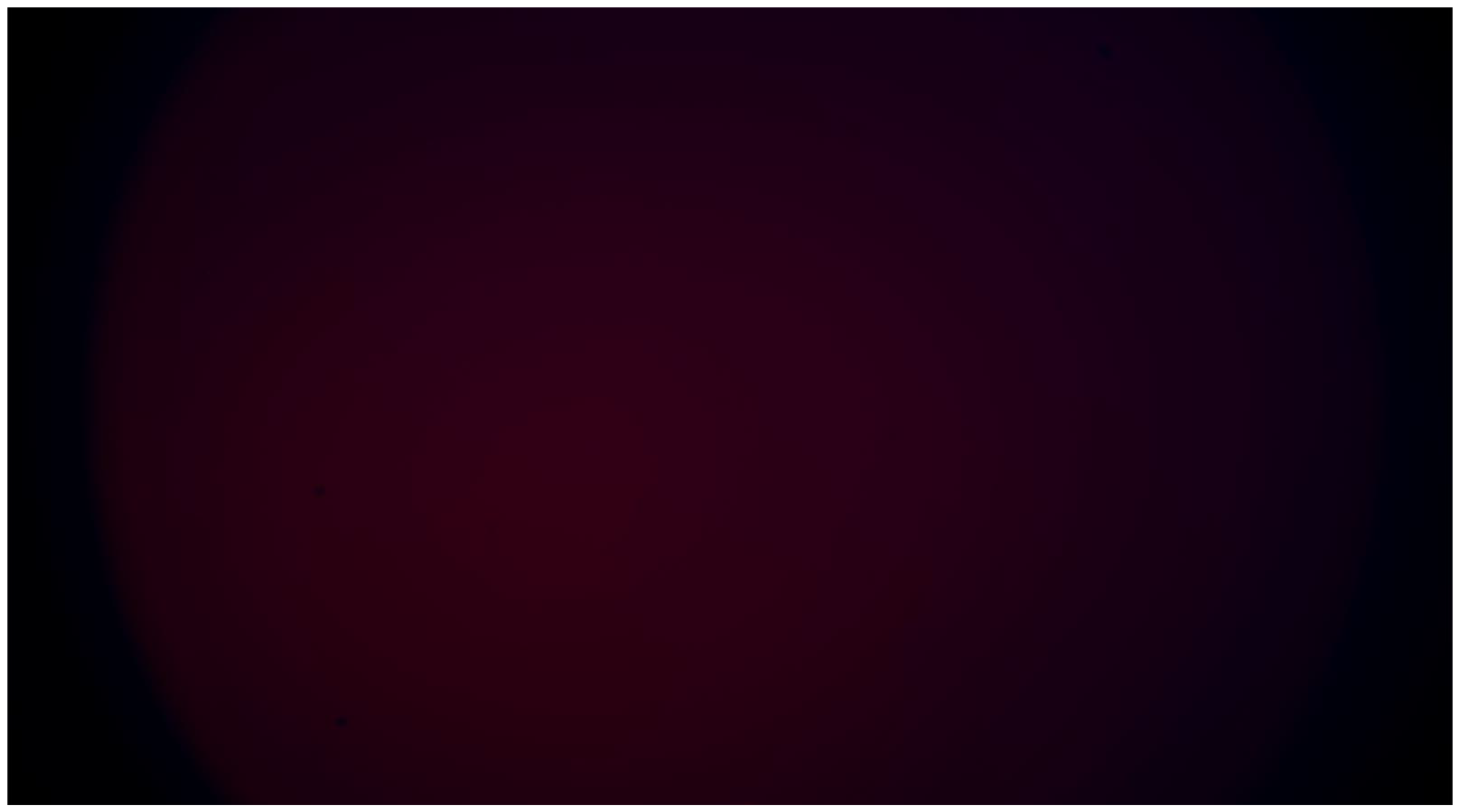
Control sample, no dyed MPs added. No fluorescence or false positives are displayed under a fluorescent microscope.

**Figure 1B:**
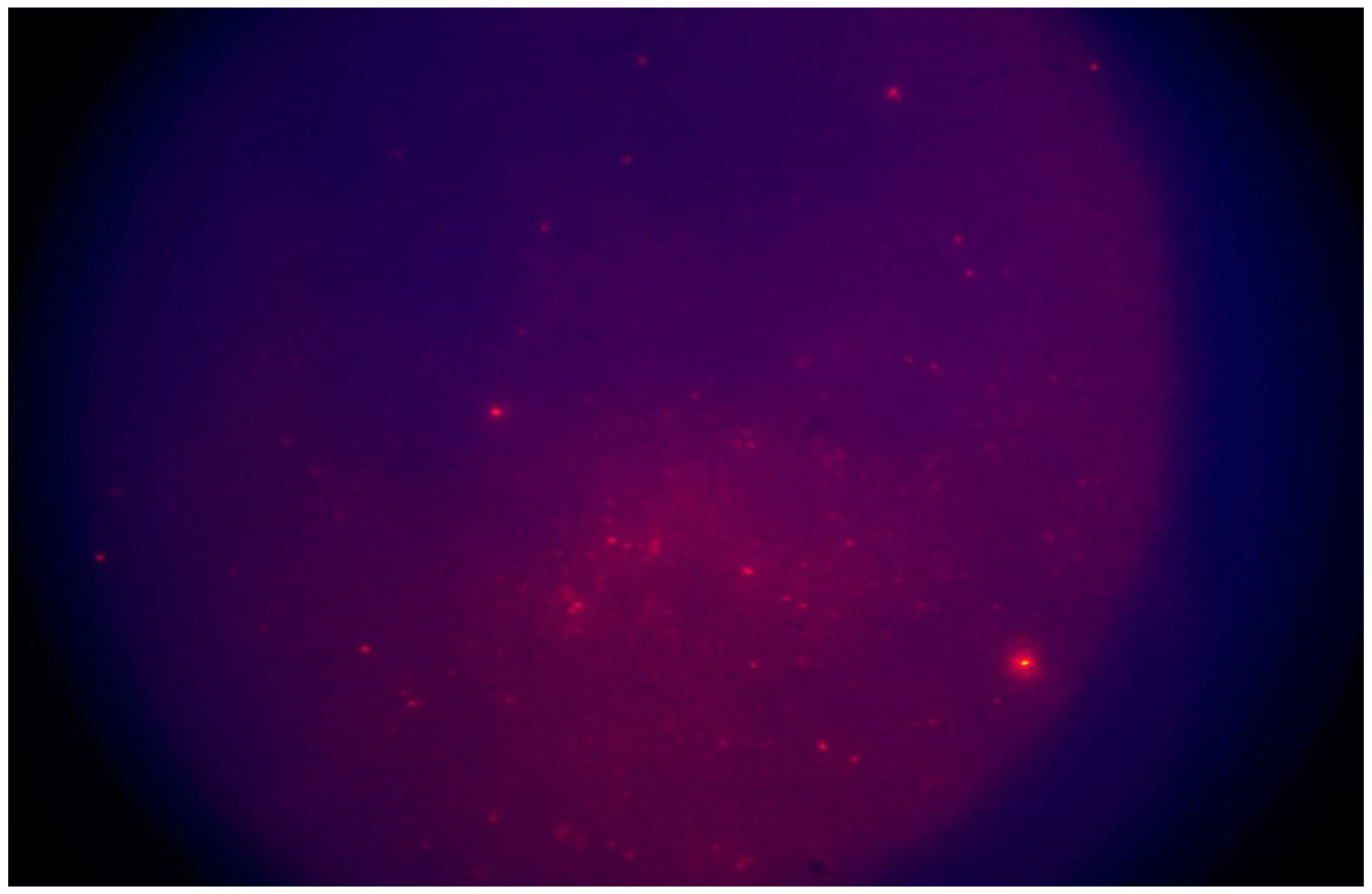
Example of MP fluorescence on filter under a fluorescent microscope

**Figure 1C:**
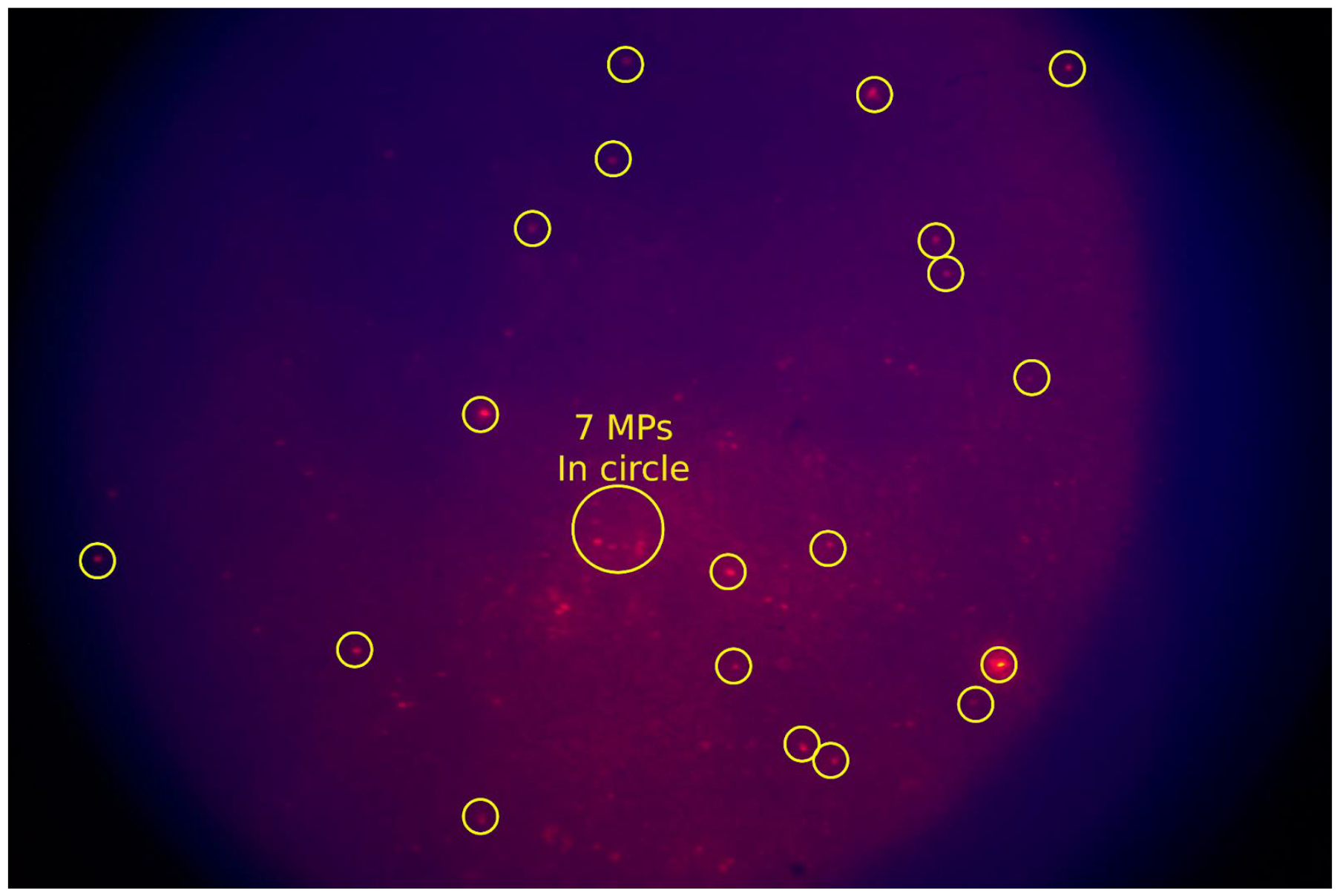
Example of MP fluorescence on filter under a fluorescent microscope. Examples of what would be counted as individual MPs are highlighted.

[Figures 1A, 1B, 1C]

Due to low sample numbers per time point, data analysis for statistically significant trends in retention over time were not performed. Nevertheless, total MP counts per filter (**Figure 2A**) or per g of wet weight (**Figure 2B**) calculated for each week of mussel collection showed a clear signal for mussel MP content at Week 1 compared to control mussels where MPs were not detectable (see Figure 1A) clear trend after an initial increase.

**Figure 2A/B:**
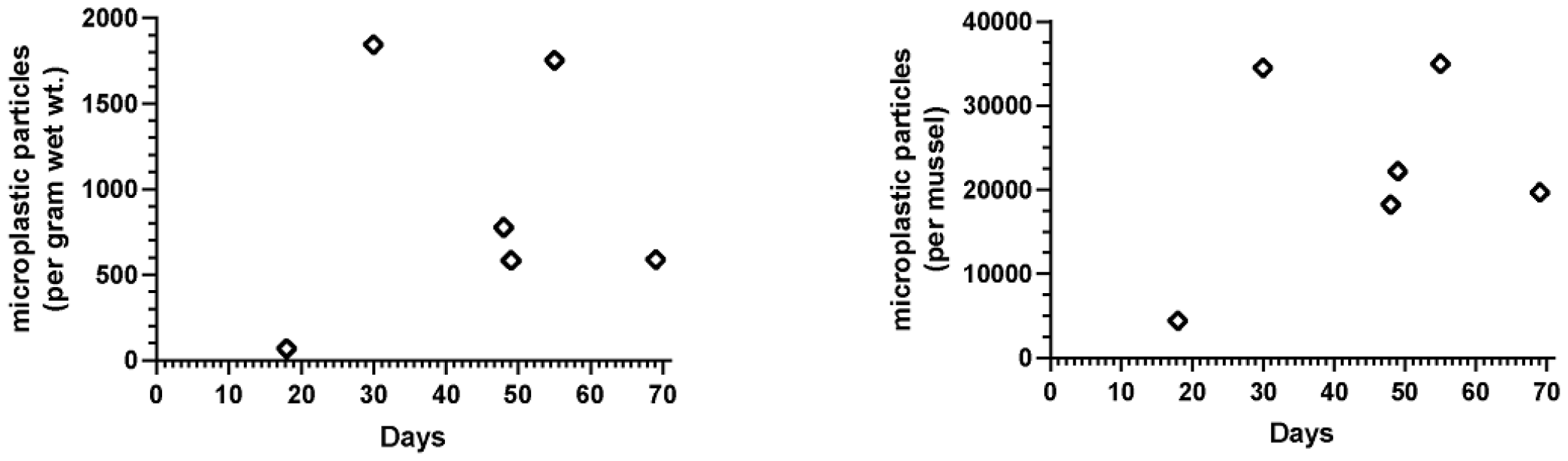
2A (left) depicts total MP counts from 1 individual mussel/week from the MP tank. No fluorescence was detected in control mussels at any of the time points. 2B (right) depicts total MP count per gram of wet tissue (without shell) for the individual mussels depicted in figure 2A.

[Figures 2A, 2B]

## Discussion

The method led to successful measurement of dyed MPs from mussel tissue. Nile Red Dye coloring was still visible following tissue digestion, storage, and sonication. In other words, the strong chemicals potassium hydroxide, nitric acid, and sodium hypochlorite successfully removed unfiltered organic matter without destroying the MPs or breaking down the dye. If indigestible body parts such as shells are removed, our method effectively eliminates the organic matter while leaving the plastic particles behind to be caught on filters.

With the promising results from the current proof-of-principle study, we are confident that the methods could be scaled to perform experiments that track California Mussel MP net uptake over time and/or test how differing MP exposures influence bioaccumulation of MPs. In addition, the method could be applied to biological samples in studies that quantify MP infiltration of several populations within an ecosystem, or several species of significance to the fisheries industries. For example, due to concerns about human MP consumption from shellfish, crustaceans, and fish of many species, the current method could be used to evaluate MP exposures in the food supply. Measuring MPs in different organisms or foods would also be useful in future studies that determine if there are associations between MP exposure levels and physiological outcomes or health perturbations. Testing the use of dye on samples post-sonication to measure natural microplastics, rather than just those added experimentally, will be an important next step.

A strength of the approach used herein was the use of mixed MPs derived from different plastics used in consumer products, thus having relevance to common waste streams in the U.S. and many other countries. Another strength of the method is the relative ease of sample processing and MP quantitation. A limitation to the study design was that analyses were limited to 1 mussel per treatment tank per week. In addition, samples from a lower dose tank (4 mg of MPs) were not available for all time points; thus, these data were not presented and no conclusions can be made related to dose-response outcomes. Despite these limitations, the primary aim to evaluate a facile, quantitative method for MPs in mussels was successful. This method should prove very useful for future studies of MPs in various experimental paradigms and in environmental samples. Shedding light on the MP consumption and retention in *Mytilus californianus* could provide new insights as to the potential roles of mussels as natural filters and bioindicators of MP pollution.

## Figures

**Figure 1A**: Example of a result from a mussel collected from the control tank (no MPs added) filter under fluorescent microscope.

**Figure 1B**: Representative illustration of MPs fluorescence signals on filter paper. Sample from a mussel treated for 1 month with the 8 mg/tank treatment.

**Figure 1C**: The same image from Figure 1B, with some examples of individual MPs circled. The labeled MP dots were counted individually from a variety of fields in order to calculate total MPs per plate (see Methods).

**Figure 2A**: Total MP counts from 1 individual mussel/week from the MP tank. Sampling began 1 week after adding the initial MP batch. No fluorescence was detected in control mussels at any of the time points. See Figure 1A.

**Figure 2B**: Total MP counts per gram of wet tissue weight (no shell) for the individual mussels depicted in Figure 2A. No fluorescence was detected in control mussels at any of the time points. See Figure 1A.

[Appendix A](https://docs.google.com/document/d/1brVoxW-fVYpgedr__6MmX9BrDcAYJdMICXsHzxawBkM/edit)

